# A Shot in the Dark: Comparative Morphology of the Bioluminescent Tube Organs in Tubeshoulders (Platytroctidae)

**DOI:** 10.1101/2025.08.26.672518

**Authors:** Emily M. Carr, Rene P. Martin, Michael J. Ghedotti, John S. Sparks

**Affiliations:** American Museum of Natural History, Department of Ichthyology, Division of Vertebrate Zoology, New York; Richard Gilder Graduate School, American Museum of Natural History, New York; University of Nebraska-Lincoln, School of Natural Resources, Nebraska; Regis University, Department of Biology, Denver

**Author notes:** These authors contributed equally to this work. These authors also contributed equally to this work (Rene P. Martin, Michael J. Ghedotti, John S. Sparks).

## Abstract

Bioluminescence, light produced by a living organism, is a key innovation in the diversification of deep-sea fishes. It is useful for a myriad of behaviors and interactions in deep sea organisms, including communication, predation, camouflage via counterillumination, and predator avoidance. In this study, we investigate the deep-sea tubeshoulders (Platytroctidae), fishes that possess a unique postcleithral tube organ associated with their shoulder girdle that excretes bioluminescent fluid, a feature that unites all members of this poorly studied family. Many tubeshoulders also possess additional bioluminescent structures and luminescent tissues, including a series of tube organs on the caudal peduncle unique to *Platytroctes apus* that are hypothesized to be similar in structure and function to the postcleithral tube organ. Herein, we present the first histological analysis of the caudal tube organs in *P. apus* and use histological methods to investigate the morphological diversity in postcleithral tube organ structure across 14 species of platytroctids, representing 10 of 13 valid genera. We show that the postcleithral tube organ generally exhibits a conserved morphology across genera and species, however, several species-specific anatomical differences are noted. In some individuals, we observe the presence of luminescent-fluid cells within the tube organ in various stages of development, which may provide evidence for inferring the type of secretory gland found in this novel bioluminescent light organ. We also show that the structure of the caudal tube organs in *P. apus* are similar to the postcleithral tube organ present in all members of Platytroctidae, likely indicating a similar luminescent fluid emission function.

## Introduction

Bioluminescence is the production of light by a living organism via a chemical reaction between luciferin, a light producing molecule, and luciferase, an enzyme [1]. Bioluminescence is phylogenetically widespread and has evolved numerous times across the tree of life [2]. This functionally diverse phenomenon is hypothesized to be used in predator avoidance, prey capture, and mating behaviors [3]. Bioluminescence in some dinoflagellates and squid species functions to deter predators, whereas other organisms like deep-sea anglerfishes (Lophiiformes: Ceratioidei) use bioluminescence to attract prey [3]. Many lanternfishes (Myctophidae) have bioluminescent light organs (photophores and secondary light organs) that are species-specific or sexually dimorphic, likely aiding in interspecific recognition [4,5]. However, ventral photophores in lanternfishes are not tied to communication, and are instead used for counterillumination, matching the intensity of downwelling light from the surface to obscure the silhouette from predators lurking below [6]. Flashlight fishes have been shown to school based entirely on bioluminescent light emitted from their subocular organs [7]. The numerous functions of marine bioluminescence are facilitated by both a diversity of light organ structures and photogenic tissues, as well as numerous mechanisms for producing and emitting light.

Bioluminescence in marine fishes is generally achieved via one of two ways. Some fishes house symbiotic bioluminescent bacteria (e.g., *Allivibrio*, *Photobacterium*, *Vibrio*) in internal structures, whereas others produce light intrinsically via the reaction of luciferin and luciferase and emit luminescence via photophores and other photogenic tissues [3]. Although intrinsic bioluminescence is widespread in marine fishes, occurring in species-rich groups such as lanternfishes (Myctophiformes) and dragonfishes (Stomiiformes), no bioluminescent fish species has been shown to synthesize their own luciferin, and all are hypothesized to obtain it via their diet [1,5,8,9].

Bioluminescent fishes also exhibit considerable variation in their light emitting structures, which can range from simple patches of bioluminescent tissue [4] to significantly more anatomically complex structures, including internal light organs, as well as light emitting photophores, chin barbels, and lures [5,10–12]. For example, ponyfishes (Leiognathidae) can control the emission of bacterially-generated bioluminescent light from a circumesophageal light organ via a complex, species-specific system of muscular lenses, clearing of the guanine-lined gas bladder, and translucent patches on the flank and other body regions [13–16]. In flashlight fishes (Anomalopidae) occlusion of subocular luminescence is achieved either via muscular rotation of the organ itself, an elastic skin shutter mechanism, or a combination of both depending on the genus [17]. The primary photophores in lanternfishes are complex in structure, containing a reflector cup and modified scale that acts as a lens to direct and possibly alter the wavelength of emitted bioluminescent light [10].

However, one type of luminescence that is considerably less common in fishes involves structures that emit, eject, or secrete bioluminescent fluid into the surrounding environment [18,19]. Secretory luminescence has independently evolved in numerous marine lineages of invertebrates and vertebrates, including annelids, cephalopods, ostracods, copepods, mysid and decapod shrimp, elasmobranchs, and teleosts [18]. Secretory luminescence is generally hypothesized to function in predator avoidance by creating a distraction or “smoke screen”, allowing light-emitting organisms to escape [20,21]. However, some ostracods also use their luminescent secretions for sexual displays where males have been shown to exhibit species-specific light flashes [22]. Other species, like the vampire squid (*Vampyroteuthis infernalis*), have viscous, mucous-like luminescent secretions that presumably stick to potential predators and mark them with light, potentially attracting additional predators [23,24]. Interestingly, most species with luminescent secretions produce their bioluminescence endogenously, and not from symbioses with luminescent bacteria [18].

Although the function of secretory luminescence is hypothesized to be relatively conserved across lineages, both the location of these light organs and the cellular mechanisms facilitating the production of the secretion are highly variable. In copepods, luminous glands can be found along the entire body (head, abdomen, and caudal furcae) and consist of single-celled sacs opening to a pore through the outer cuticle [20], whereas ostracods possess a glandular light organ on their upper lip, expelling luminescent fluid via muscle contractions [25]. Vampire squid (*Vampyroteuthis infernalis*) secrete bioluminescent mucus from each arm tip [24,26], while the cuttlefish *Hetrroteuthis dispar* secretes luminescent fluid from its siphon, sometimes mixing it with ink [27]. Multiple species of shrimp (e.g., Oplophoridae, *Heterocarpus, Thalassocaris, Glyphus*) expel luminescent fluid from their mouth, which is likely the result of a hepatic process [28,29].

In vertebrates, luminous secretions are limited to a single genus of marine elasmobranch (*Mollisquama*) [18], which comprises two morphologically similar species, and a single family of bony fishes (Platytroctidae) [30]. In the American Pocket Shark (*Mollisquama mississippiensis*) the light organ is comprised of pectoral pockets and is hypothesized to secrete luminescent fluid via epidermal holocrine glands, whereby expulsion is triggered via movement of the pectoral fin [18]. However, the mechanism of secretory luminescence remains poorly understood in Platytroctidae.

Platytroctidae is a morphologically diverse family comprising 40 species arrayed within 13 genera of globally distributed (except the Mediterranean Sea) deep-sea fishes ranging from mesopelagic to bathypelagic depths (200-4000 m)^25^. Members of the family, commonly called tubeshoulders, are united by the presence of an apomorphic postcleithral tube organ, a unique sac-like internal structure situated beneath the cleithrum that can emit bioluminescent fluid into the surrounding environment via an external tube^25^ (Fig. 1). This strategy is hypothesized to be used for the same function as other secretory luminescent taxa, either to escape predation or ward off predators^25^. As a result, members of this family have been informally referred to as “luminous cloud-throwers” [31]. Many platytroctids also possess additional discrete light organs embedded in ventral regions of their epidermis thought to be used in ventral counterillumination [19], as well as morphologically distinct patches of luminescent tissue on several different body regions that vary considerably among genera. In one species, *Platytroctes apus,* a series of openings on the dorsal and ventral margins of the caudal peduncle leading to darkly pigmented chambers have been mentioned by previous authors, although these structures have not been studied in detail^25^. Our knowledge regarding the internal structure and function of the postcleithral tube organ in Platytroctidae, as well as the caudal organs present in *P. apus*, remains limited.

**Fig 1.**
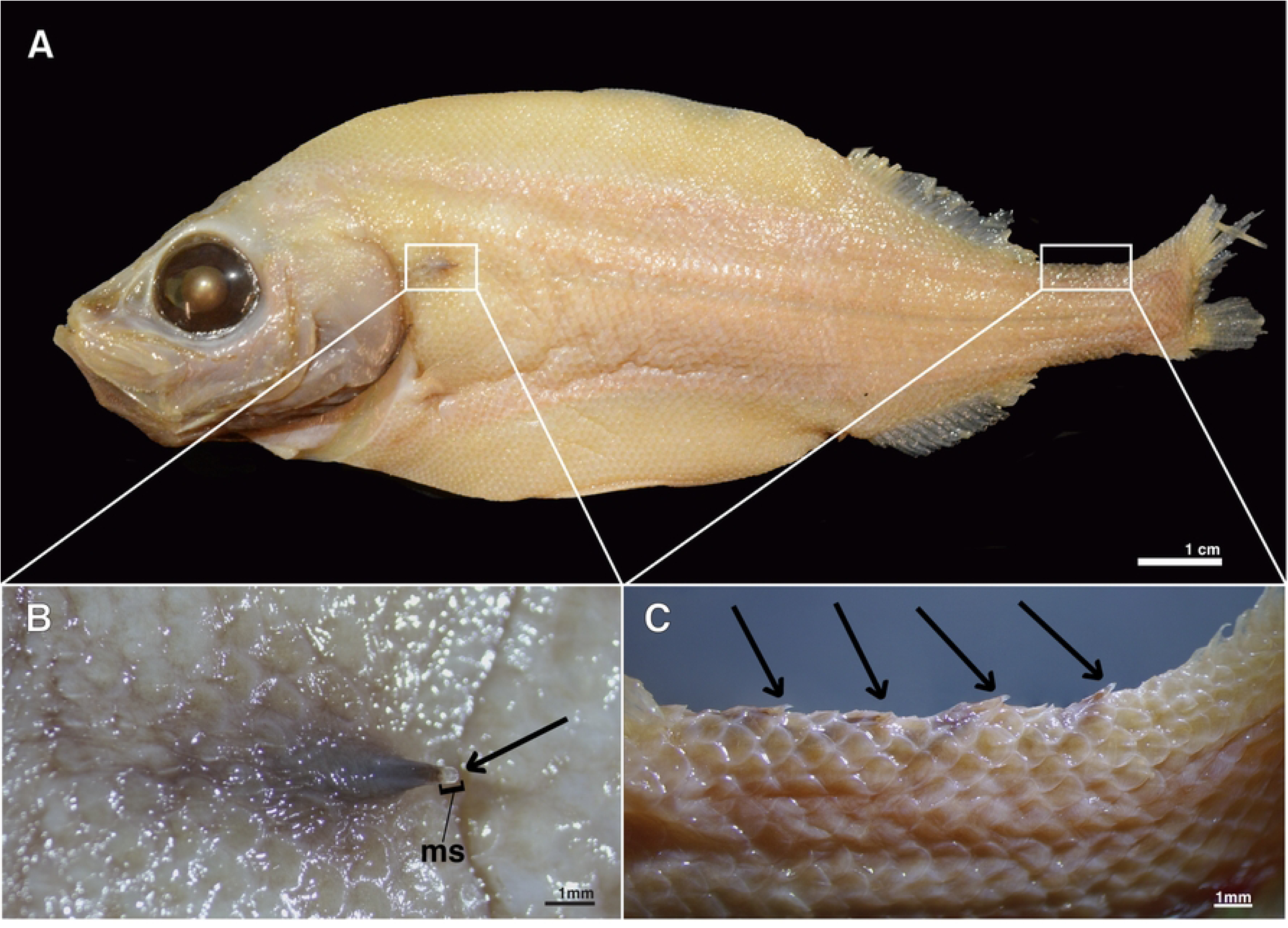
External anatomy of the postcleithral and caudal tube organ. (A) *Platytroctes apus* (SIO 55-244, 136.4 mm SL) showing location of postcleithral tube organ (left white box) and caudal tube organs (right white box). (B) Postcleithral tube organ in *P. apus* (SIO 77-54) showing terminal opening and modified scale (ms). (C) Caudal tube organs on dorsal margin of caudal peduncle in *P. apus* (SIO 55-244). Arrows indicate the terminal opening of the postcleithral and caudal tube organs.

Beebe [32] documented the reduction of the external projection of the tube organ over ontogeny and described a modified scale that supports the tube in one platytroctid species, *Mentodus rostratus* [32]. Histological analyses of the tube organ are limited to a single study that examined only one species, *Sagamichthys schnakenbecki* [21], and we know very little regarding the composition of the bioluminescent fluid of the tube organ or its glandular basis. Overall, the postcleithral tube organ in platytroctids remains extremely understudied and virtually no variation in this organ has yet been documented among platytroctid species.

In this study we describe the morphology and ultrastructure of the postcleithral tube organ across Platytroctidae using histological techniques. The goals of this study are to 1) describe tube organ ultrastructure and compare across platytroctids, and 2) document and formally describe similar tube-like organs that occur on the upper and lower margins of the caudal peduncle in *Platytroctes apus*. This study is critical in advancing our knowledge of platytroctid tube organs as a luminescence ejection system, an extremely rare phenomenon in fishes. We present the first taxonomically comprehensive comparative analysis of the tube organ across 10 of 13 recognized platytroctid genera using high-resolution color images of histological sections. We also provide the first histological sections and detailed description of the structure of the caudal tube organs that are unique to *P. apus*.

## Methods

All specimens sectioned for histological analysis were formalin-fixed and alcohol-preserved (75% ethanol or 50% isopropanol) museum specimens from multiple institutions (Table 1). Gross anatomy of the tube organs (e.g., the presence of modified scales) was examined using a Nikon SMZ800N stereo microscope. The tube organ (right side) and surrounding tissue was dissected for histological preparation in 14 platytroctid species representing 10 of the 13 total genera (Fig. 1B; Table 1). Two of the dorsal caudal tube organs were also dissected for histological sectioning in *Platytroctes apus* (n=1; Fig. 1C). The presence of the caudal tube organ was determined externally by examining the caudal peduncle of all species using a dissecting Nikon SMZ800N stereo microscope. Many platytroctid species are rarely collected, and as a result, are represented by few specimens in museum collections. Thus, dissections on additional individuals were often not possible. Specimens examined for presence of caudal tube organs in addition to specimens used for histological analyses are listed in the supplementary materials (Fig. S1).

**Table 1.** Platytroctid species examined for histological analysis of tube organ structure in this study.

| Genus | Species | Catalog number | Standard Length (mm) | Modified Scale in Postcleithral Tube Organ | Caudal Tube Organ |
| --- | --- | --- | --- | --- | --- |
| *Barbantus* | *curvifrons* | MCZ 60746 | 77.2 | Present | Absent |
| *Holtbyrnia* | *latifrons* | SIO 80-110 | 174.4 | Present | Absent |
|  | *macrops* | LACM 30432-26 | 21.9 | Absent | Absent |
|  | *melanocephala* | LACM 31065-9 | 34.9 | Present | Absent |
| *Maulisia* | *microlepis* | MCZ 168334 | 121.6 | Present | Absent |
| *Mentodus* | *facilis* | LACM 9706-42 | 46.2 | Present | Absent |
| *Mirorictus* | *taningi* | SIO 14-136 | 118.1 | Present | Absent |
| *Normichthys* | *operosus* | MCZ 163242 | 129.7 | Present | Absent |
|  | *yahganorum* | SIO 61-45 | 42.8 | Present | Absent |
| *Platytroctes* | *apus* | SIO 77-54 | 130.2 | Present | Present |
|  | *mirus* | MCZ 138114 | 101.8 | Present | Absent |
| *Sagamichthys* | *abei* | SIO 18-4 | 209.2 | Present | Absent |
| *Searsia* | *koefoedi* | MCZ 168583 | 137.3 | Present | Absent |
| *Searsioides* | *multispinus* | SIO 61-32 | 101.9 | Present | Absent |

The presence of the caudal tube organ was determined externally by examining the caudal peduncle of all species using a dissecting Nikon SMZ800N stereo microscope.

Tissues were dehydrated through a stepwise series of increasing concentrations of EtOH to 100% and incubated in xylene for clearing. Tissue was subsequently embedded in paraffin wax and sectioned laterally through the tube organ at 5-10 μm intervals using a Leica HistoCore MULTICUT rotary microtome. Sections were mounted on charged slides with distilled water and dried at 50°C for 24 hours. Slides were then stained with Mason’s Trichrome following the protocol from Ghedotti et al. [12]. Stained sections were mounted in Canada balsam and imaged using a Nikon Eclipse 50*i* compound microscope equipped with an Excelis 4K UHD microscope camera. Images used in figures were prepared by increasing brightness, eliminating erroneous debris caused by the mounting process, and by combining photographs into composite images using Adobe Photoshop. Institutional abbreviations of specimens used for histological analysis in this study are: Natural History Museum of Los Angeles County (LACM), Museum of Comparative Zoology, Harvard University (MCZ), and Scripps Institution of Oceanography, University of California, San Diego (SIO).

## Results

### Postcleithral Tube Organ

We observed overall similar morphology in the postcleithral tube organ across all species examined, including *Barbantus curvifrons, Holtbyrnia latifrons, H. macrops, H. melanocephala, Maulisia microlepis, Mentodus facilis, Mirorictus taningi, Normichthys operosus, N. yahganorum, Platytroctes apus, P. mirus, Sagamichthys abei, Searsia koefoedi,* and *Searsioides multispinus* (Figs. 2-4, Table 1). The postcleithral tube organ is located ventral to the lateral line, posterior to the opercular series, and ventroposterior to the cleithrum (Fig. 1A). The external tip (= opening) of the tube is oriented posteriorly and is often associated with a modified scale (Fig. 1B) in all species examined except for *H. macrops* (Figs. 1, 2-4). When present, this modified scale (ms) is curved into a conical shape and extends internally (i.e., inside the lumen of the tube for about 1/3 to 1/2 of its length; Fig. 2) to support the external opening of the tube organ (Figs. 2-4). The tube organ (t) lies within the dermis (d) below the scales (s) and epidermis, with the lumen of the usually scale-supported tube extending anteroventrally from the external opening (tip) of the organ and located adjacent and superficial to the hypaxial musculature (m) of the body wall. Within the tube organ, a dense pigment layer (pl) of melanophores lines the wall of the lumen, becoming denser apically, nearer the opening to the external environment (Fig. 2).

**Fig 2.**
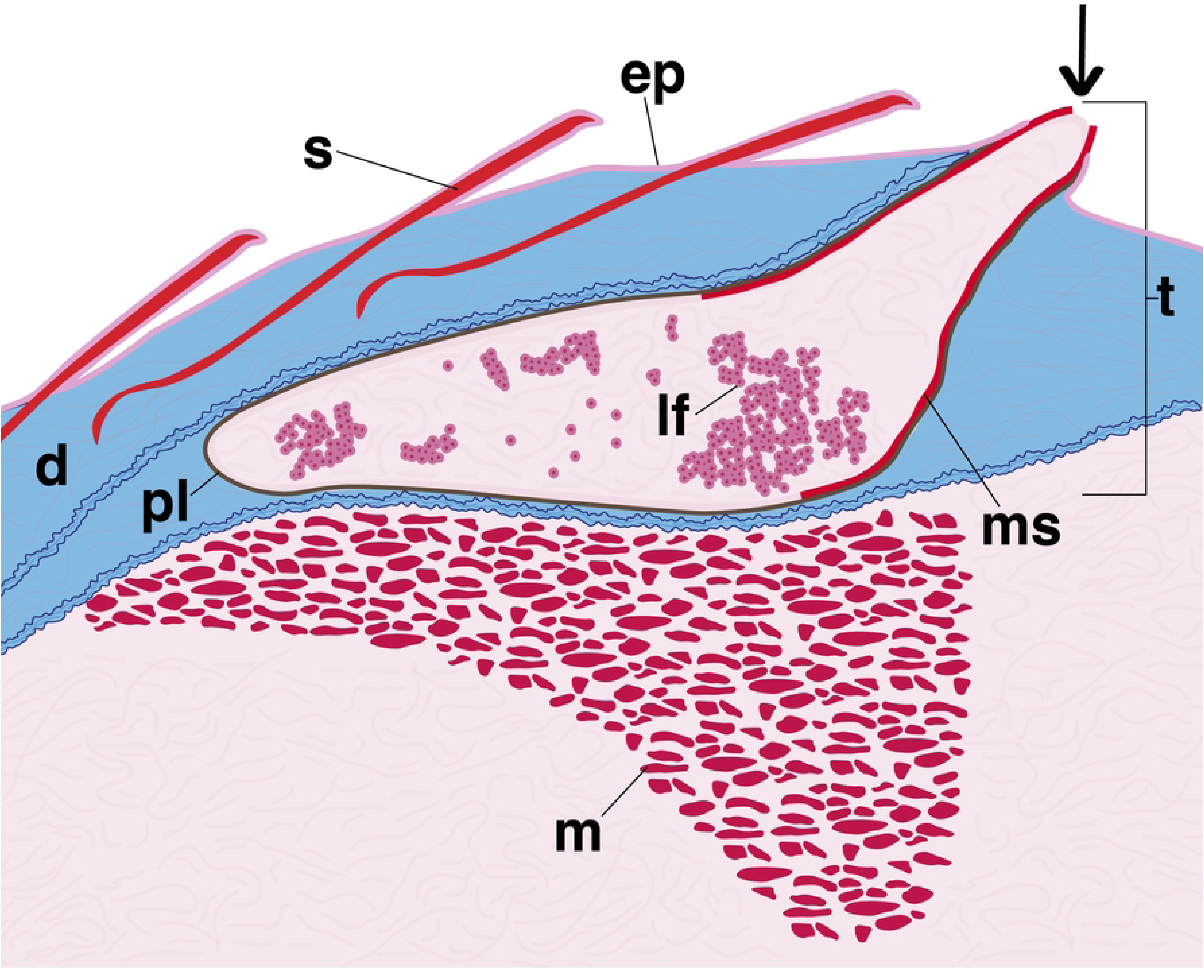
Generalized illustration showing internal postcleithral tube organ anatomy. Abbreviations: scale (s, red), epithelium (ep, dark pink), dermis (d, blue), tube organ (t), pigment layer (pl, brown), modified scale (ms, red), luminescent fluid (lf, pink), muscle (m, red). Arrow indicates terminal tube opening, which is directed posteriorly.

The lumen of the postcleithral tube organ often contains spherical luminescent-fluid cells (lf) (Figs. 2-4). These cells have darkly stained nuclei with surrounding granules that stain red (Fig. 3A and C). In some species, the granules of the luminescent-fluid cells are lightly stained, and the nucleus is clear (Fig. 3B). The diameter of the luminescent-fluid cells ranges from 13-32 μm, with an average diameter of 19.4 μm. In most, but not all species, smaller cells are also present within the dermis surrounding the lumen of the tube organ (*) (Fig. 3C-D), often associated with the pigment layer (pl). These cells have very limited cytoplasm that is lightly stained relative to the fully developed luminescent-fluid cells (lf), but the nuclei remain darkly stained (Fig. 3C-D). Although the size of these cell clumps (*) (Fig. 3C-D) varies greatly among species, the component cells have a consistent diameter of 3-6 μm, with an average cell size of 4.5 μm. A third cell type, blue cells (bc), are also found in the lumen of the tube (Fig. 3C). Blue cells (bc) have a darkly stained nucleus and an unstained or lightly blue stained cytoplasm, ranging in diameter from 8-12 μm (Fig. 3C). The postcleithral tube organ is also highly vascularized, and we observed the presence of numerous oblong red blood cells (rb; 8-12 μm) that are contained within blood vessels (v; Fig. 3C).

**Fig 3.**
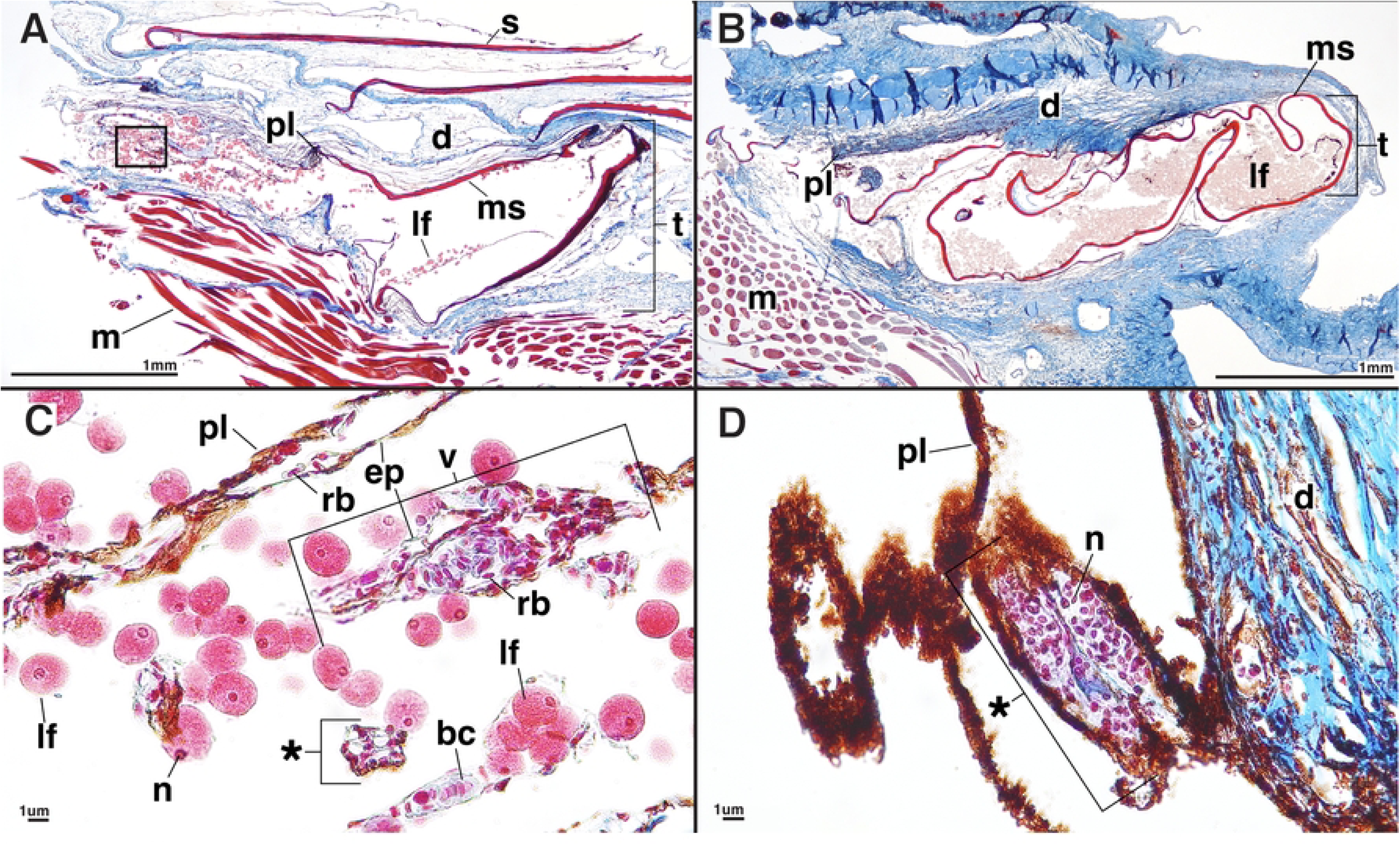
Longitudinal sections through the postcleithral tube organ. (A) *Mirorictus taningi* (SIO 14-136) and (B) *Searsia koefoedi* (MCZ 168583). Potential developing luminescent-fluid cells (*) in (C) *Mirorictus taningi* (SIO 14-136) and (D) *Platytroctes apus* (SIO 77-54). Anterior is to the left in all figure panels, such that the posteriorly-directed terminal opening of the tube organ is to the right. Abbreviations: scale (s), dermis (d), tube organ (t), pigment layer (pl), modified scale (ms), luminescent fluid (lf), muscle (m), nucleus (n), blue cells (bc), red blood cells (rb), blood vessel (v), epithelium (ep). Scale bar = 10um.

**Fig 4.**
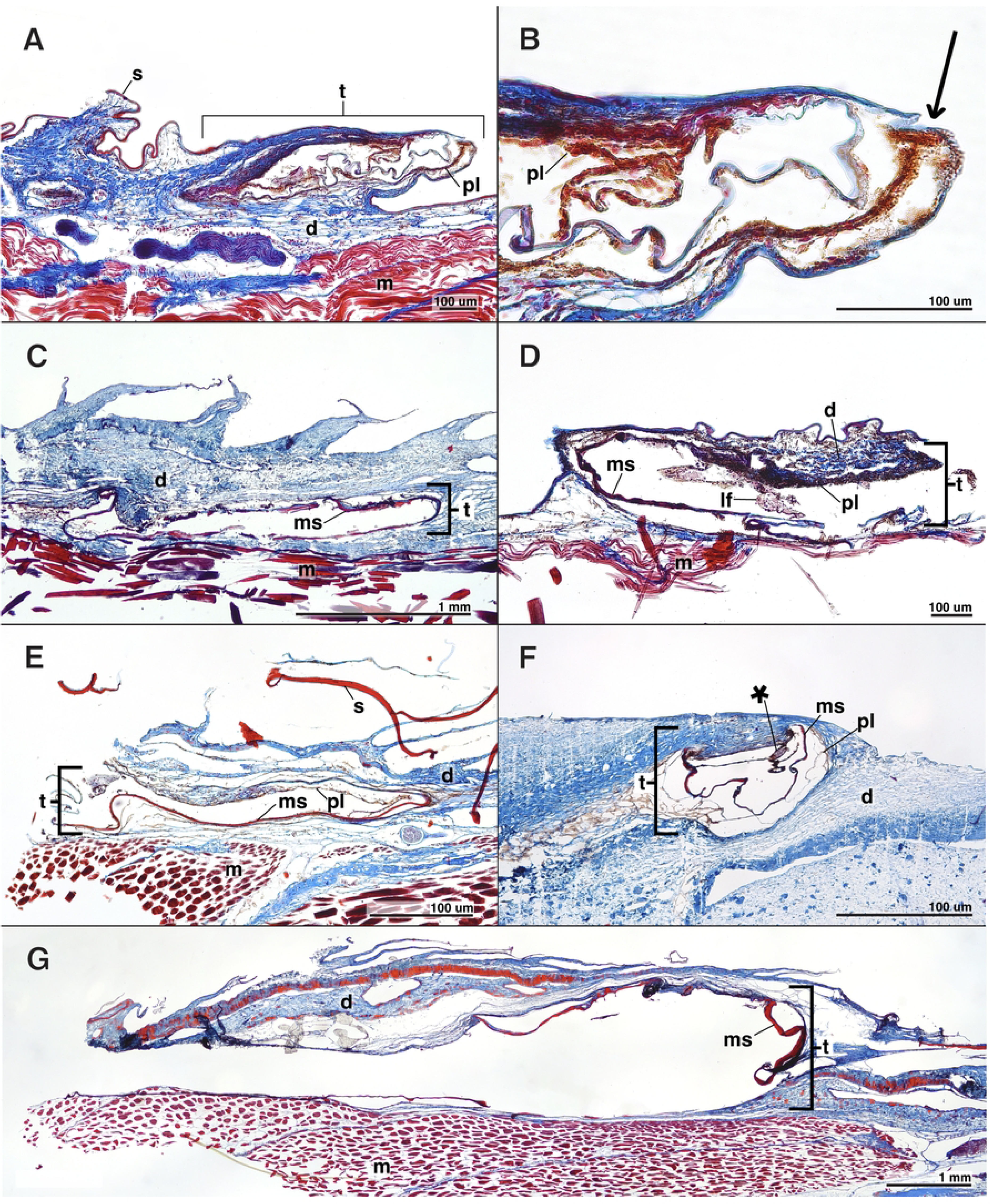
Longitudinal sections of the postcleithral tube organ. (A and B) *Normichthys yahganorum* (SIO 61-45), (C) *Normichthys operosus* (MCZ 163242), (D) *Holtbyrnia melanocephala* (LACM 31065-9), (E) *Mentodus facilis* (LACM 9706-42), (F) *Platytroctes apus* (SIO 77-54), and (G) *Sagamichthys abei* (SIO 18-4). Anterior is to the left in all figure panels, such that the posteriorly-directed terminal opening of the tube organ is to the right. Abbreviations: scale (s), dermis (d), tube organ (t), pigment layer (pl), modified scale (ms), luminescent fluid (lf), muscle (m), blood vessel (v), potential developing luminescent-fluid cells (*). Arrow in panel B points towards terminal tube opening.

### Caudal Peduncle Tube Organs

In *Platytroctes apus* we note the presence of tube organs on the caudal peduncle which have a similar morphology to the postcleithral tube organ (Fig. 1C). These organs are darkly pigmented, arranged in series, and number from two to five distinct darkly pigmented (i.e., black) tube-like structures on both the dorsal and ventral margins of the caudal peduncle depending on the specimen examined (Figs. 1C and 5). Each caudal tube organ has an associated modified scale (ms) that is concavely curled and slightly elevated compared to adjacent scales (Figs. 1C and 5). We note the presence of numerous luminescent-fluid cells (lf) in each of the caudal light organs, ranging in size from 14-26 μm (Fig. 5A). We also observe smaller cells (*) in the walls of the caudal light organs, ranging in size from 4-8 μm (Fig. 5B). We find that caudal tube organs are lacking entirely (i.e., they are not present on either the dorsal or ventral margins of the caudal peduncle) in *P. mirus*, the only other currently recognized species in the genus *Platytroctes*, as well as in all other platytroctid species.

**Fig 5.**
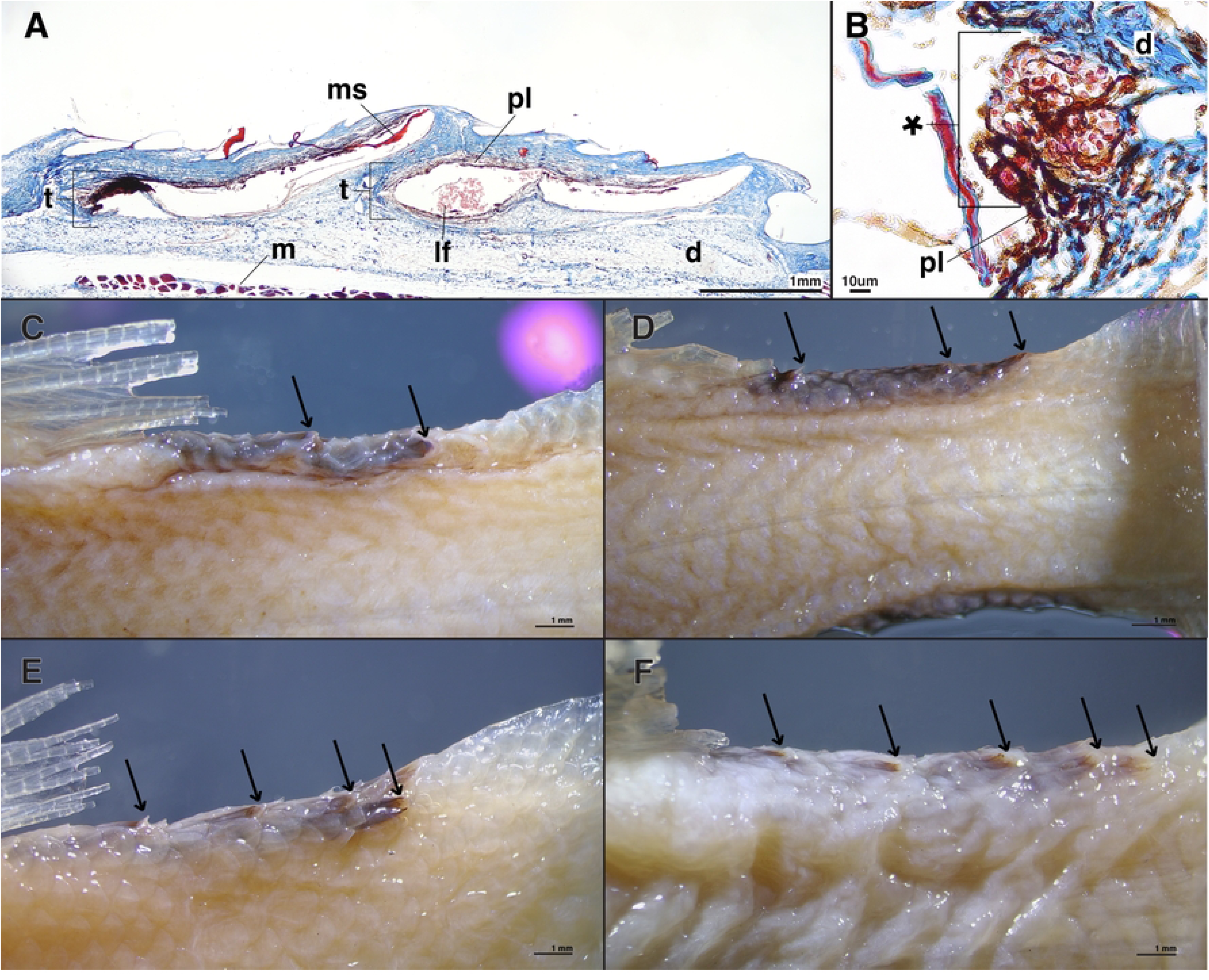
Longitudinal histological sections of the caudal tube organ in P. apus (SIO 77-54). (A) Two dorsal caudal tube organs in lateral view and (B) potential developing luminescent-fluid cells (*) in the lumen of a caudal tube organ in *P. apus*. Variation in the number of dorsal caudal tube organs in *Platytroctes apus*, showing (C) two (USNM 206894), (D) three (USNM 206894), (E) four (USNM 206893), and (F) five (USNM 201649) caudal tube organs. Arrows point towards terminal caudal tube opening. Note: Ventral tube organs on the caudal peduncle in *P. apus* also vary in number. Anterior is to the left in all figure panels, such that the posteriorly-directed terminal openings of the caudal tube organs are located to the right. Abbreviations: dermis (d), tube organ (t), pigment layer (pl), modified scale (ms), luminescent fluid (lf), muscle (m).

## Discussion

In this study we describe the morphology and ultrastructure of the postcleithral tube organ in 10 of the 13 genera of Platytroctidae and provide the first detailed description of the caudal tube organs unique to *Platytroctes apus*. We find that the overall structure of the postcleithral tube organ is generally conserved across Platytroctidae. However, a few species-specific anatomical differences are noted and described below. We observe a dark pigment layer along the internal surface of the tube organ in all species investigated (Figs. 3 and 4). This aligns with previous observations of the dark internal lining of the tube organ [32]. However, we find that this pigment layer is much thicker on the internal apical surface of tube organ lumen, closer to the external environment (Figs. 3 and 4). We also observe that some species exhibit a significantly thicker pigment layer than others. *Holtbyrnia melanocephala* (Fig. 4D) has an extremely thick pigment layer in the dermis lining the postcleithral tube, whereas the pigment layer is thin and reduced in *Mirorictus taningi* (Fig. 3A). Pigment layer thickness is likely not related to habitat depth, as mesopelagic genera (e.g. *Holtbyrnia*, *Searsia*) [30] have pigment layers of varying thickness. Instead, thicker pigment layers in the postcleithral tube organ are observed in species with darker overall body pigmentation (e.g. *H. melanocephala,* Fig. 4D). A similar pigment layer is seen in the light organ of the only other vertebrate species known to emit bioluminescent fluid, the American Pocket Shark (*Mollisquama mississippiensis*), which are hypothesized to expel bioluminescent fluid via the movement of the closely associated pectoral fin [18]. This layer of pigmentation likely functions to prevent light from the bioluminescent fluid from being visible to the external environment while it is within the tube organ prior to secretion.

A modified scale is found to support the tube organ in all but one species that we examined (Figs. 1B, 3A-B, and 4). The absence of a modified supporting scale in *Holtbyrnia macrops* may be due to accidental removal during collection via abrasive forces, given that our specimen of *H. macrops* is relatively small (21.9 mm SL) and lacks any observable body scales. The modified scale is hypothesized to provide structure and stability to the tube organ, particularly distally, near its opening to the environment (Fig. 2) [31], although it likely also functions to direct the luminescent fluid during expulsion [19]. At the terminal, external tip of the tube, the modified scale is folded into a funnel-like shape (Fig. 1B), which likely directs the luminescent fluid during expulsion^28^. For example, in live *Searsia koefoedi*, applying gentle external pressure to the shoulder region resulted in a jet of luminescent fluid being ejected from the tube organ up to 25 cm in the air [33]. However, Herring [33] also states that recently deceased specimens of *S. koefoedi* in air responded to the same stimulus by slowly releasing the luminescent fluid onto the flank of the organism [33], akin to observations of luminescent secretions in recently deceased *Sagamichthys schnakenbecki* [19]. This may suggest a mechanism of muscular control of the expulsion of luminescent fluid, perhaps by contraction of the adjacent hypaxial muscle layer (Figs. 3 and 4).

### Tube Organ Cell Types / Comparison to Nicol [21] (1958)

We find similar cell types within the postcleithral tube organ to those reported by Nicol [21], the only histological study of the postcleithral tube organ in platytroctids to date. Despite only investigating one species, *Sagamichthys schnakenbecki*, Nicol [21] found three potentially luminescent cell types associated with the tube organ: red-stained spherical cells in the lumen (8-40 μm), violet-stained cells in the walls and lumen of the tube (5um), and blue-stained cells in the lumen (6-8 μm). Our histological observations reveal a large number of luminescent-fluid cells (lf) of similar size (13-32 μm) to Nicol’s “red cells” [21] in the lumen of the tube organ (Figs. 3 and 4). We also find smaller cells (3-6 μm) that are stained dark violet, and that are often associated with the pigment layer (Fig. 3C and D). These cells (*) are likely the “violet cells” observed by Nicol [21]. We also observe slightly larger cells that have an unstained or very light blue cytoplasm with a dark violet nucleus, likely the “blue cells” (bc) reported by Nicol [21] (Fig. 3C) that are noticeably rounder and contain more cytoplasm.

Nicol [21] hypothesized that the “violet cells” were generative tissue that pass into the lumen of the gland and transition to the “blue cells”, which develop into larger, luminous “red cells” [i.e., luminescent fluid (lf) cells]. However, Nicol [21] was hesitant regarding this hypothesis, as no such exocrine luminous secretions are known to be dermally derived [18,34]. The platytroctid postcleithral tube organ lacks a prominent epithelial layer with secretory cells (e.g., goblet cells) or an apical layer from a stratified epithelium that could be the source of the cellular luminescent discharge. It is possible that the small violet cells (*) (Fig. 3C and D) we observe could be clumps of epithelium. However, we are uncertain whether these rarely observed cells represent true epithelial cells, or if they are even present in a sufficient quantity to potentially support the number of luminescent-fluid cells observed. Interestingly, we do find a thin layer of epithelium associated with the numerous blood vessels that are present in the collagen-rich connective tissue near the walls of the tube organ (Fig. 3C). We usually find red blood cells (8-12 μm) within these vessels as well (Fig. 3C). Thus, whereas our observed cell types are consistent with Nicol’s [21] observations, the hypothesized mechanism of a dermally derived exocrine gland is unlikely. Instead, it seems more plausible that the luminescent-fluid cells, and the potential precursor cell types, originate outside the tube organ, perhaps in the proximate hematopoietic head kidney. However, further studies are needed to determine whether a direct connection exists between the postcleithral tube organ and head kidney.

### Caudal Peduncle Tube Organs

In a single species, *Platytroctes apus*, we observe the presence of tube-like organs of a similar morphology to the postcleithral tube organ on the caudal peduncle. These organs are arranged in a median series and number from two to five darkly pigmented distinct tube-like organs on both the dorsal and ventral margins of the caudal peduncle depending on the specimen examined (Figs. 1C and 5). Based on our histological analyses, we find that the caudal tube organs in *P. apus* are structurally similar to the postcleithral tube organ common to all members of Platytroctidae. Each tube-like caudal organ has a modified scale supporting the external opening of the tube organ, a darkly pigmented lining, and all three cell types of the developing luminescent fluid as described above (Fig. 5). However, the modified supporting scale in the caudal tube organs is folded into a U-shape and oriented away from the body (Fig. 1C and 5), unlike the modified supporting scale of the postcleithral tube organ, which folds tightly inward towards the body (Fig. 1B). As a result, the caudal tube organs lack a distinct tube tip, and instead resemble more of a terminal scoop. Based on the overall similar anatomy of the postcleithral and caudal tube organs, and the presence of the same luminescent-fluid cells in both types of organ, we hypothesize that the caudal tube organs of *P. apus* can emit bioluminescent fluid and that they function similarly to the postcleithral tube organ [35]. *Platytroctes mirus*, the only other currently recognized species in the genus *Platytroctes*, lacks caudal tube-like organs entirely, as we did not observe them on either the dorsal or ventral margins of the caudal peduncle.

## Conclusion

In this study we investigate the structure of the postcleithral tube organ across 14 species of platytroctids, representing 10 of 13 total genera (Table 1; Figs. 1B, 2-4). We provide the first histological analysis and detailed description of the unique caudal tube organs found only in *Platytroctes apus*, showing that these organs are similar in structure to the postcleithral tube organ (Figs. 1C, 5). We find that the postcleithral tube organ is a highly vascularized sac-like structure with a modified supporting scale, an internal apically distributed layer of dark pigment lining the tube lumen, and with no continuous epithelial layer lining the lumen of the organ (Figs. 2-4). We also observe three cell types in agreement with the original findings of Nicol [21] in both the postcleithral tube organ of all species examined, as well as the caudal tube organs of *P. apus*: clumps of smaller cells (*) in the tube organ wall, poorly stained “blue cells” in the lumen, and numerous larger luminescent-fluid cells also in the lumen (Fig. 3).

Whereas we agree with Nicol’s [21] observations regarding cell types present in the tube organs, we propose an alternative hypothesis regarding the origin of the luminescent fluid in the postcleithral tube organ. Due to the lack of a layer of epithelium and a high degree of vascularization, the bioluminescent fluid in the platytroctid postcleithral tube organ may originate via hematopoietic processes in the proximate head kidney. Although we have not yet been able to determine whether there is a direct connection between the postcleithral tube organ and the head kidney, we note that the head kidney and lumen of the postcleithral tube organ are in close proximity. In order to test this hypothesis, further studies are needed to understand the metabolic processes leading to the production of functional luminescent cells found in the platytroctid postcleithral tube organ and determine whether a direct connection exists between the postcleithral tube organ and head kidney. Regardless, luminescent cell production appears to be the result of a unique metabolic mechanism in tubeshoulders and unlike anything documented to date in teleost fishes.

## Acknowledgements

We would like to thank B. Frable (SIO), G. Lauder and A. Williston (MCZ), and B. Ludt and T. Clardy (LACM) for the loan of specimens in their care and for permission to dissect the tube organ for histological analysis. We also thank P. Rask Moller (ZMUC), P. Bartsch and E. Abel (ZMB), and T. Weddehage (ZMH) for their hospitality and access to collections in their care.

## Supporting Information Captions

**S1 Fig.** Specimens examined for presence of caudal tube organs in addition to specimens used for histological analyses listed in Table 1.

## Notes

### Competing Interest Statement

The authors have declared no competing interest.

## References

1. Davis MP, Sparks JS, Smith WL. Repeated and Widespread Evolution of Bioluminescence in Marine Fishes. Thuesen EV, editor. PLOS ONE. 2016;11: e0155154. doi:10.1371/journal.pone.0155154

2. Lau ES, Oakley TH. Multi-level convergence of complex traits and the evolution of bioluminescence. Biol Rev. 2021;96: 673–691. doi:10.1111/brv.12672

3. Haddock SHD, Moline MA, Case JF. Bioluminescence in the Sea. 2009.

4. Davis MP, Holcroft NI, Wiley EO, Sparks JS, Leo Smith W. Species-specific bioluminescence facilitates speciation in the deep sea. Mar Biol. 2014;161: 1139–1148. doi:10.1007/s00227-014-2406-x

5. Martin RP, Davis MP, Smith WL. The impact of evolutionary trade-offs among bioluminescent organs and body shape in the deep sea: A case study on lanternfishes. Deep Sea Res Part Oceanogr Res Pap. 2022;184: 103769. doi:10.1016/j.dsr.2022.103769

6. Paitio J, Yano D, Muneyama E, Takei S, Asada H, Iwasaka M, et al. Reflector of the body photophore in lanternfish is mechanistically tuned to project the biochemical emission in photocytes for counterillumination. Biochem Biophys Res Commun. 2020;521: 821–826. doi:10.1016/j.bbrc.2019.10.197

7. Gruber DF, Phillips BT, O’Brien R, Boominathan V, Veeraraghavan A, Vasan G, et al. Bioluminescent flashes drive nighttime schooling behavior and synchronized swimming dynamics in flashlight fish. PLOS ONE. 2019;14: e0219852. doi:10.1371/journal.pone.0219852

8. Bessho-Uehara M. Kleptoprotein bioluminescence: *Parapriacanthus* fish obtain luciferase from ostracod prey. Sci Adv. 2020;6: 4942.

9. Mirza J, Oba Y, Mirza J, Oba Y. Semi-Intrinsic Luminescence in Marine Organisms. Bioluminescence - Technology and Biology. IntechOpen; 2021. doi:10.5772/intechopen.99369

10. Martin RP, Carr EM, Sparks JS. Variation in lanternfish (Myctophidae) photophore structure: A comprehensive comparative analysis. PLOS ONE. 2024;19: e0310976. doi:10.1371/journal.pone.0310976

11. Ghedotti MJ, Smith WL, Davis MP. The first evidence of intrinsic epidermal bioluminescence within ray-finned fishes in the linebelly swallower *Pseudoscopelus sagamianus* (Chiasmodontidae). J Fish Biol. 2019;95: 1540–1543. doi:10.1111/jfb.14179

12. Ghedotti MJ, DeKay HM, Maile AJ, Smith WL, Davis MP. Anatomy and evolution of bioluminescent organs in the slimeheads (Teleostei: Trachichthyidae). J Morphol. 2021;282: 820–832. doi:10.1002/jmor.21349

13. McFall-Ngai M, Montgomery MK. The Anatomy and Morphology of the Adult Bacterial Light Organ of *Euprymna scolopes* Berry (Cephalopoda:Sepiolidae). Biol Bull. 1990;179: 332–339. doi:10.2307/1542325

14. Sparks JS, Dunlap PV, Smith WL. Evolution and diversification of a sexually dimorphic luminescent system in ponyfishes (Teleostei: Leiognathidae), including diagnoses for two new genera. Cladistics. 2005;21: 305–327. doi:10.1111/j.1096-0031.2005.00067.x

15. Sparks JS, Dunlap PV. A Clade of Non-Sexually Dimorphic Ponyfishes (Teleostei: Perciformes: Leiognathidae): Phylogeny, Taxonomy, and Description of a New Species. Am Mus Novit. 2004;2004: 1–21. doi:10.1206/0003-0082(2004)459%3C0001:ACONDP%3E2.0.CO;2

16. Chakrabarty P, Davis MP, Smith WL, Baldwin ZH, Sparks JS. Is sexual selection driving diversification of the bioluminescent ponyfishes (Teleostei: Leiognathidae)? Mol Ecol. 2011;20: 2818–2834. doi:10.1111/j.1365-294X.2011.05112.x

17. Johnson GD, Rosenblatt RH. Mechanisms of light organ occlusion in flashlight fishes, family Anomalopidae (Teleostei: Beryciformes), and the evolution of the group. Zool J Linn Soc. 1988;94: 65–96. doi:10.1111/j.1096-3642.1988.tb00882.x

18. Claes JM, Delroisse J, Grace MA, Doosey MH, Duchatelet L, Mallefet J. Histological evidence for secretory bioluminescence from pectoral pockets of the American Pocket Shark (*Mollisquama mississippiensis*). Sci Rep. 2020;10: 18762. doi:10.1038/s41598-020-75656-8

19. Poulsen JY. New observations and ontogenetic transformation of photogenic tissues in the tubeshoulder *Sagamichthys schnakenbecki* (Platytroctidae, Alepocephaliformes). J Fish Biol. 2019;94: 62–76. doi:10.1111/jfb.13857

20. Clarke GL, Conover RJ, David CN, Nicol J a. C. Comparative studies of luminescence in copepods and other pelagic marine animals. J Mar Biol Assoc U K. 1962;42: 541–564. doi:10.1017/S0025315400054254

21. Nicol J a. C. Observations on luminescence in pelagic animals. J Mar Biol Assoc U K. 1958;37: 705–752. doi:10.1017/S0025315400005749

22. Gerrish GA, Morin JG. Living in sympatry via differentiation in time, space and display characters of courtship behaviors of bioluminescent marine ostracods. Mar Biol. 2016;163: 190. doi:10.1007/s00227-016-2960-5

23. Mirza JD, Migotto ÁE, Yampolsky IV, de Moraes GV, Tsarkova AS, Oliveira AG. *Chaetopterus variopedatus* Bioluminescence: A Review of Light Emission within a Species Complex. Photochem Photobiol. 2020;96: 768–778. doi:10.1111/php.13221

24. Robison BH, Reisenbichler KR, Hunt JC, Haddock SHD. Light Production by the Arm Tips of the Deep-Sea Cephalopod *Vampyroteuthis infernalis*. Biol Bull. 2003;205: 102–109. doi:10.2307/1543231

25. Huvard AL. Ultrastructure of the light organ and immunocytochemical localization of luciferase in luminescent marine ostracods (Crustacea: Ostracoda: Cypridinidae). J Morphol. 1993;218: 181–193. doi:10.1002/jmor.1052180207

26. Hoving HJT, Robison BH. Vampire squid: detritivores in the oxygen minimum zone. Proc R Soc B Biol Sci. 2012;279: 4559–4567. doi:10.1098/rspb.2012.1357

27. Dilly PN, Herring PJ. The light organ and ink sac of *Heteroteuthis dispar* (Mollusca: Cephalopoda). J Zool. 1978;186: 47–59. doi:10.1111/j.1469-7998.1978.tb03356.x

28. Chan BK, Lin I-C, Shih T-W, Chan T-Y. Bioluminescent emissions of the deep-water pandalid shrimp, *Heterocarpus sibogae* De Man, 1917 (Decapoda, Caridea, Pandalidae) under laboratory conditions. Crustaceana. 2008;81: 341.

29. Herring PJ. Bioluminescence in the Crustacea. J Crustac Biol. 1985;5: 557–573.

30. Matsui T, Rosenblatt RH. Review of the Deep-Sea Fish Family Platytroctidae (Pisces: Salmoniformes). 1987.

31. Tucker DW. Report on the fishes collected by SY“ Rosaura” in the North and Central Atlantic, 1937-38. Part I. Families Carcharhinidae, Torpedinidae, Rosauridae (nov.), Salmonidae, Alepocephalidae, Searsidae, Clupeidae. Bull Brit MusNat Hist. 1954;2: 163–214.

32. Beebe W. Deep-sea fishes of the Bermuda Oceanographic Expeditions Family Alepocephalidae. Zoologica. 1933;16: 15–93.

33. Herring PJ. Bioluminescence of searsid fishes. J Mar Biol Assoc U K. 1972;52: 879–887. doi:10.1017/S0025315400040613

34. Genten F, Terwinghe E, Danguy A. Atlas of fish histology. CRC Press; 2009.

35. Sazonov YI. I. Light. Morphology and significance of the luminous organs in alepocephaloid fishes. In: Deep Sea and Extreme Shallow-water Habitats: Affinities and Adaptations. Biosyst Ecol. 1996;11: 151–163.

